# Ovarian support cell in vitro maturation (OSC-IVM) results in healthy murine live births with no evidence of reprotoxicology in a multigenerational study

**DOI:** 10.1101/2024.04.04.588122

**Authors:** Maria Marchante, Ferran Barrachina, Enric Mestres, Monica Acacio, Kathryn S Potts, Sabrina Piechota, Bruna Paulsen, Alexander D Noblett, Alexandra B Figueroa, Nuno Costa-Borges, Christian C Kramme

## Abstract

**Study question:** Does application of human stem cell-derived ovarian support cells (OSCs) for *in vitro* maturation (IVM) have a safe reproductive toxicity profile?

**Summary answer:** The use of OSC-IVM co-culture improves blastocyst formation in a mouse model and results in healthy live births with no evidence of reprotoxicity.

**What is known already:** Abbreviated stimulation to obtain immature oocytes combined with a successful IVM offers a promising alternative to traditional *in vitro* fertilization, reducing hormonal doses and making IVF shorter and safer. Recently, we developed an OSC platform derived from human induced pluripotent stem cells (hiPSCs) that replicate dynamic ovarian function *in vitro*, enhancing human oocyte maturation and yielding an improved blastocyst formation rate compared to commercial IVM options. However, the reproductive toxicity profile, commonly assessed via murine multigenerational models, for OSC-IVM remains unknown.

**Study design, size, duration:** A total of 70 B6/CBA 6–8-week-old stimulated female mice were used in this study to collect immature mouse oocytes (n=2,025) at the germinal vesicle (GV) stage. Half of these oocytes were retrieved denuded (denuded oocytes condition, n= 930), while the remaining oocytes were kept with the cumulus cells (COCs condition, n= 1,095) to simulate the two possible dispositions of oocytes during clinical practice. Oocytes from each condition, denuded oocytes and COCs, were randomly assigned to either commercially available traditional IVM media (MediCult-IVM^TM^, Origio) group (control group) or the same traditional IVM media supplemented with human OSCs (Fertilo^TM^, Gameto Inc.) to form the OSC-IVM group (test group).

**Participants/materials, setting, methods:** Oocytes from each condition, denuded oocytes and COCs, were subjected to *in vitro* culture for 18-20 hours. After IVM, metaphase II (M2) oocytes were inseminated by intracytoplasmic sperm injection (ICSI) and cultured to assess blastocyst formation *in vitro*. Embryos that reached the blastocyst stage on day five were vitrified using Kitazato’s protocol in preparation for embryo transfers. A group of M2 oocytes and blastocyst embryos were employed for quality analyses by immunofluorescence.

Vitrified blastocysts were warmed and transferred to pseudopregnant females (4-5 embryos per uterine horn), evaluating the F1 offspring. Pup characteristics were tracked, including weight, length, sex ratio, and physiology. Weekly monitoring assessed mouse behavior and development. At reproductive age, select F1 mice were outbred to wildtype mice to produce the F2 generation, analyzing live births, sex ratio, morphology, and behavior across groups. Moreover, hormonal and organ histological analyses were performed in F1 mice to further explore the overall health of the progeny.

**Main results and the role of chance:** In contrast to findings in humans, in mice OSC-IVM generally led to a decreased maturation rate compared to Traditional-IVM (68.6% ± 14.1% versus 80.9% ± 5.9%, p=0.0101). Subsequent embryo culture yielded significantly different fertilization rates between the four groups (p=0.0055). Specifically, OSC-IVM with COCs significantly differed from Traditional-IVM with denuded oocytes (89.5 ± 10.5 versus 96.5 ± 4.8, p=0.0098). There were no differences in the cleavage rates (p=0.7547). However, there was a significant distinction in the blastocyst formation (p=0.0068), wherein OSC-IVM with COCs showed a greater formation rate compared to Traditional-IVM for both denuded oocytes and COCs (56.1% ± 19.2% versus 41.5% ± 15.9% and 38.0% ± 16.2%; p=0.0408, and p=0.0063). Spindle morphology analysis demonstrated normal spindle morphology in denuded oocytes and COCs under both Traditional-IVM and OSC-IVM. Moreover, embryo analysis showed no significant difference in inner cell mass count (p=0.1550).

Following embryo transfers, analysis of live births showed no significant distinctions between groups regarding delivery, sex ratio, pup length, developmental and behavioral abnormalities, hormonal values or histopathological anomalies in the F1 generation. Evaluation of the F2 generation also showed no significant differences in live births, sex ratio, or developmental/behavioral abnormalities between groups, further validating the absence of long-term implications and transgenerational effects derived from OSC-IVM culture.

**Limitations, reasons for caution:** Although this study was conducted in compliance with European Medicines Agency (EMA) ICH E6 (R2) Good clinical practice scientific guidelines to demonstrate the OSC safety, human clinical studies evaluating in vivo and live birth outcomes are necessary to corroborate the findings of this study.

**Wider implications of the findings:** This study provides evidence of the safety of using the OSC-IVM system, as evidenced by the lack of adverse effects on *in vitro* embryo development post OSC-IVM and on the health and fertility of offspring across successive generations *in vivo*.

**Trial registration number:** N/A

## Introduction

In recent years, *in vitro* maturation (IVM) of oocytes has emerged as a promising technique, offering novel possibilities for fertility treatment (De Vos et al., 2021; Gilchrist and Smitz, 2023). Unlike conventional *in vitro* fertilization (IVF) methods, which rely on controlled ovarian stimulation (COS) through the administration of exogenous hormones at high doses (Macklon et al., 2006), IVM enables the retrieval of immature oocytes from unstimulated or minimally stimulated ovaries, allowing for their maturation outside of the body. By reducing the doses of gonadotropins required for ovarian stimulation, this approach not only mitigates the potential adverse effects associated with high hormonal dosage, but also minimizes the risk of ovarian hyperstimulation syndrome (OHSS), a serious complication of conventional COS (Mostinckx et al., 2024; Zhang et al., 2016). Moreover, it eliminates the discomfort and inconvenience associated with hormone injections, further enhancing the appeal of IVM. Consequently, this innovative technique not only shortens the duration of treatment, but also substantially reduces medication costs, thereby rendering IVF more affordable. Additionally, IVM may offer a more cost-effective and accessible option compared to conventional IVF ( Souza et al., 2023), particularly for patients with conditions such as polycystic ovary syndrome (PCOS) or those susceptible to ovarian complications (Cela et al., 2018; De Vos et al., 2021; Gilchrist and Smitz, 2023; Mahajan, 2015; Schirmer et al., 2020).

Reproductive clinics worldwide are increasingly incorporating IVM into their treatment protocols, leading to the birth of 5,000-6,000 infants through this technique (Das and Son, 2023; Yang and Chian, 2017). However, ongoing research and advancements in IVM protocols are necessary to enhance success rates and broaden their clinical applicability. Current IVM protocols predominantly rely on supplementing cell culture media with growth factors, small molecules, and hormones known to influence oocyte meiotic progression (Akin et al., 2021; Braam et al., 2019; De Vos et al., 2011; Fadini et al., 2009; Guzman et al., 2012; Ma et al., 2020; Mochini et al., 2011; Mohsenzadeh et al., 2022; Piechota et al., 2023; Pongsuthirak et al., 2015; Sanchez et al., 2019; Sanchez et al., 2017; Shu-Chi et al., 2006; Vuong et al., 2020a, 2020b). However, these protocols have yielded various outcomes concerning oocyte maturation and embryo formation rates (Gilchrist et al., 2024), often attributed to challenges associated with the synchronization of nuclear-cytoplasmic maturation and the heterogeneity of oocyte disposition at retrieval (Ahmad et al., 2023). Importantly, it is well known that cumulus encasement of the oocyte is critically important to influence the rate and quality of oocyte maturation *in vitro*, but that often during the retrieval and manipulation prior to IVM, mechanical shearing may result in oocytes with or without the cumulus enclosure. Hence, there is the need for innovative IVM approaches that leverage advanced formulations tailored to mimic the complex microenvironment of oocyte follicle *in vivo*, aimed at enhancing the oocyte quality and competence.

Recently, our group has developed a novel strategy, termed OSC-IVM, aimed at improving the efficiency of IVM. OSC-IVM involves an oocyte co-culture system with ovarian support cells (OSCs) derived from human induced pluripotent stem cells (hiPSCs) that reproduce the dynamic ovarian function *in vitro*. Leveraging their functional resemblance to human granulosa cells, Fertilo OSCs enhance the nuclear and cytoplasmic maturation of human oocytes *in vitro* by supplementation into traditional IVM media. Consequently, OSC-IVM significantly improves human oocyte maturation rates and subsequent embryo development, yielding an improved blastocyst formation rate compared to traditional IVM options (Piechota et al., 2023; Giovannini et al 2023).

Despite the numerous advantages of the novel OSC-IVM system, a systematic analysis of the approach’s reproductive toxicity profile has not yet been performed, a key necessity for its clinical implementation. A multigenerational mouse reproductive model is a regulatory gold standard for therapeutics and provides an evaluation to ensure the safety and effectiveness of the technique. Therefore, this study aimed to provide robust evidence regarding the absence of reproductive toxicity associated with OSC-IVM by comparison of murine *in vitro* embryo development, health, and the fertility of offspring across successive generations *in vivo* after treatment of murine oocytes with and without cumulus enclosure using traditional IVM media with and without human OSC supplementation.

## Materials and Methods

### Study Design

This study was conducted as a pivotal toxicology assessment of the recently described OSC-IVM method to assess the impact of co-culturing human ovarian support cells (OSCs) with immature oocytes (Piechota et al. 2023). To achieve this, fresh immature mouse oocytes from the B6/CBA hybrid strain were collected at the germinal vesicle (GV) stage from stimulated females. The oocytes were allocated into two different groups based on their morphology upon retrieval: a subset of retrieved oocytes was naturally denuded of their cumulus cells (“denuded” oocyte group), while the remaining oocytes retained their surrounding cumulus cells, constituting the cumulus-oocyte complex (COC) group. Subsequently, immature oocytes from each condition were subdivided and cultured under two conditions: in OSC-IVM composed of traditional IVM medium (MediCult-IVM, Origio) supplemented with human OSCs (Fertilo, Gameto Inc.) (test group) or in traditional IVM media (Medicult-IVM, Origio) without OSCs (control group). Importantly, this study evaluated the use and safety of OSC-IVM for both cumulus enclosed oocytes (COCs) and cumulus denuded oocytes, to model all possible human applications for IVM treatment.

The present study was divided into two phases: The first phase (*in vitro*) aimed to evaluate oocyte maturation, oocyte quality via meiotic spindle analysis, and *in vitro* embryo development comparing both OSC-IVM versus Traditional IVM systems. The second phase of this study (*in vivo*) focused on studying the long-term effects of the OSC-IVM co-culture system on reproductive potential *in vivo,* involving a comprehensive assessment of multigenerational health (F1 and F2). The study design is presented in Figure 1.

**Figure 1.**
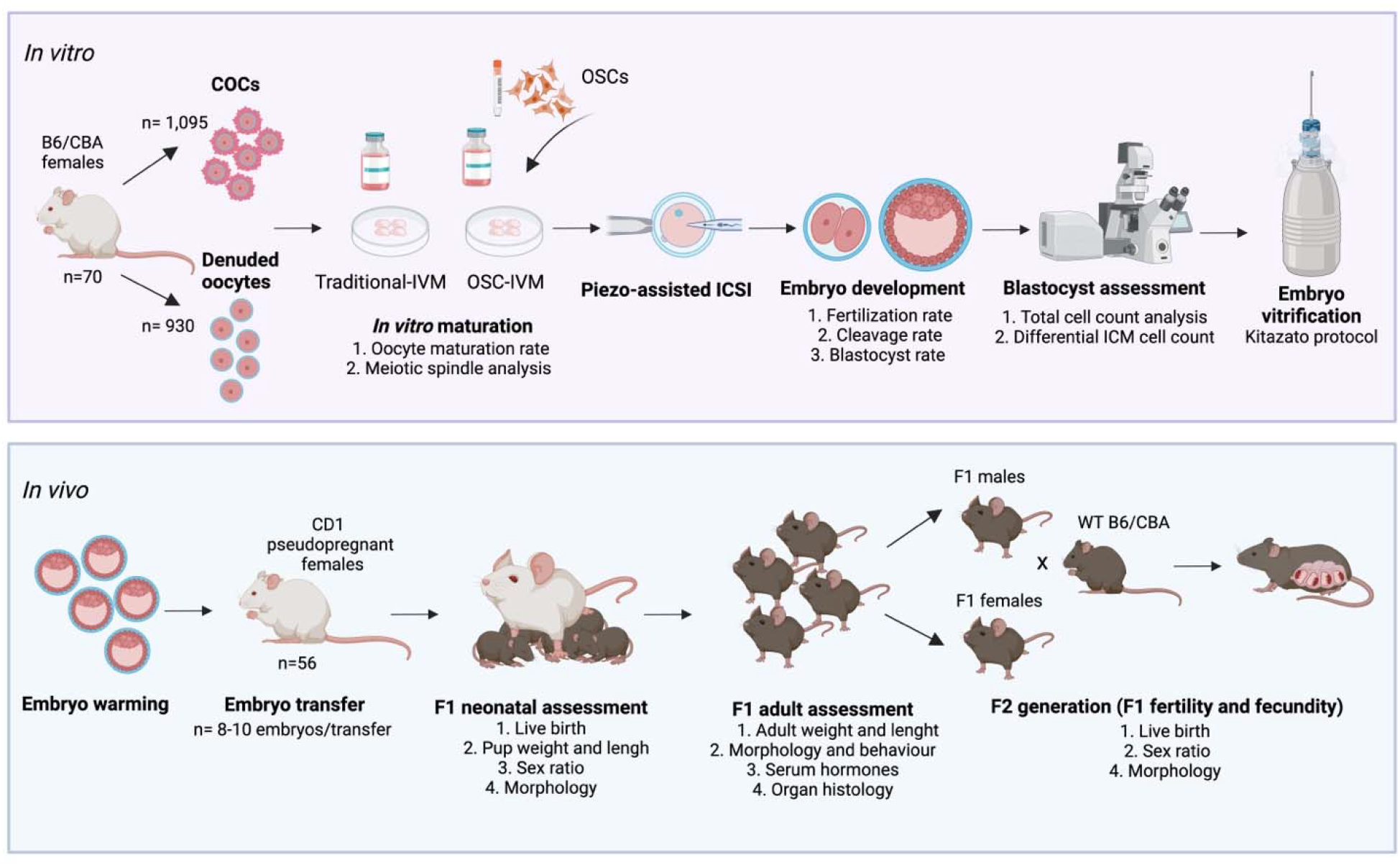
Experimental design. A. *In vitro* study. The initial *in vitro* study involved oocyte collection, *in vitro* maturation (IVM) with Traditional-IVM media (Control) or OSC-IVM containing ovarian support cells (OSCs) for 18-20 hours, fertilization using intracytoplasmic sperm injection (ICSI), embryo culture until blastocyst stage, and embryo vitrification. B. *In vivo* study. The *in vivo* section of the study encompassed embryo warming and embryo transfer, followed by the evaluation of the development and multigenerational health (F1 and F2 generation assessment). Illustration created with BioRender.com.

### Animals

All animal care and procedures were conducted according to protocols approved by the Ethics Committee on Animal Research of the Science Park of Barcelona (PCB), Spain, and in compliance with European Medicines Agency (EMA) ICH E6 (R2) Good clinical practice scientific guidelines at Embryotools (Barcelona, Spain).

Hybrid (B6/CBA), outbred CD1 females, and male mice from the same genetic backgrounds were purchased from Janvier Laboratories (Le Genest-Saint-Isle, France). Upon arrival, all mice were quarantined and acclimated to the PCB Animal facility (PRAAL) for approximately one week before use. Mice were housed with a 12-hour light/dark cycle (lights on at 7:00 A.M.) with ad libitum access to food and water. All procedures involving mice (e.g., animal handling, and administration of hormones for superovulation of females or matting), were conducted at the PRAAL Animal Facility, while procedures involving embryo manipulation (e.g oocytes collection, embryo culture up to blastocyst, blastocyst cell counts, vitrification/warming of blastocysts and embryo selection for transfer) were performed at Embryotools’ laboratories.

### Cell and media preparation

Ovarian support cells (OSCs) were generated from a single line of human induced pluripotent stem cells (hiPSCs) using overexpression of the transcription factors *GATA4*, *NR5A1* and *RUNX2*, as previously described (Pierson Smela et al., 2023, Piechota et al. 2023). OSCs are defined as a pure population of CD82+, FOXL2+ mural granulosa-like cells, and are known to be steroid and growth factor producing in response to follicle stimulating hormone (FSH) and androstenedione (A4) stimulation. The OSCs were produced in multiple batches and cryopreserved in vials of 120,000-150,000 live cells and stored in liquid nitrogen in CryoStor CS10 Cell Freezing Medium (StemCell Technologies). The morning of oocyte retrieval, cryopreserved OSCs were thawed for 2-3 minutes at 37°C (in a heated bead or water bath), resuspended in supplemented IVM media and washed twice using centrifugation and pelleting to remove residual cryoprotectant. Supplemented IVM media for OSC-IVM consisted of IVM media (Medicult, Origio) supplemented with 75 mIU/ml of recombinant human FSH (Millipore), 100 mIU/ml recombinant human chorionic gonadotropin (hCG) (Sigma), 500 ng/ml of A4 (Sigma), 1 µg/mL of doxycycline (StemCell Tech), and 10 mg/ml of human serum albumin (Sage). Androstenedione is supplemented into the media to recapitulate the role of theca cells and to provide a substrate for estradiol (E2) production by the OSCs. Supplemented IVM media equilibrated overnight at 37.3°C, 6% CO_2_ and 7% O_2_ under mineral oil (Hypure oil Heavy, Kitazato) was used for final resuspension. OSCs were then plated in suspension in non-adherent culture dishes at a concentration of 100,000 OSCs per 100 µl, as in our previous preclinical human studies (Piechota et al., 2023). The IVM condition containing OSCs (test article) in supplemented IVM media is referred to as OSC-IVM. A control condition only containing IVM media (Medicult, Origio), supplemented with 75 mIU/ml of recombinant FSH (Millipore), 100 mIU/ml recombinant hCG (Sigma), and 10 mg/ml of human serum albumin (Sage), was also included in this study, and is referred to as Traditional-IVM. Dishes containing the supplemented media droplets were prepared on the day prior to the oocyte retrieval, and were equilibrated overnight in a dry incubator at 37.3°C, 6% CO_2_ and 7% O_2_ under mineral oil (Hypure oil Heavy, Kitazato).

### Oocyte collection and IVM co-culture

Fresh immature mouse oocytes (n=2,025) from the B6/CBA hybrid strain were collected at the germinal vesicle (GV) stage from the ovaries of 6–8-week-old stimulated (7.5 IU of pregnant mare serum gonadotropin (PMSG) administered 48h before collection) female mice (n=70). The collected oocytes were allocated into the “denuded” (n=930) or “COC” (n=1,095) groups based on their morphology immediately after retrieval. Immature oocytes were initially cultured in LAG media (pre-IVM, Cooper Surgical) at 37 °C for 2 hours in a dry multi-drawer incubator (AD-3100 cube, Astec) with 6% COD and 7% OD. Subsequently, the oocytes were randomized into the two IVM conditions being tested, the Control group (Traditional-IVM) and the Test group (OSC-IVM), for 18-20 hours. Oocytes were distributed in the different groups as follows: Traditional-IVM with denuded oocytes (n=466 oocytes), OSC-IVM with denuded oocytes (n=464 oocytes), Traditional-IVM with COCs (n=553 oocytes), OSC-IVM with COCs (n=542 oocytes).

### Evaluation of oocyte *in vitro* maturation

After 18-20 hours of IVM culture, oocytes were removed from the IVM dishes and underwent assessment of oocyte maturation rate. The COCs and denuded oocytes were moved through two 50µl wash droplets to remove residual OSCs and IVM components. The assessment of the maturation rate of the groups containing COCs required a stripping step to remove the surrounding cumulus and corona cells via hyaluronidase treatment (Sigma-Aldrich) prior to oocyte evaluation. Oocyte maturation rate was assessed under an inverted microscope (IX73, Olympus), and oocytes were classified according to the following criteria: GV—presence of a germinal vesicle (GV), typically containing a single nucleolus within the oocyte; M1—absence of a GV within the oocyte and absence of a polar body (PB) in the perivitelline space (PVS) between the oocyte and the zona pellucida (ZP); M2—absence of a GV within the oocyte and presence of a PB in the PVS between the oocyte and the ZP. M2 oocytes were kept in culture medium under oil (Hypure oil Heavy, Kitazato) at 37°C and 6% COD, and 7% OD concentration until ICSI.

### Oocyte meiotic spindle analysis

For analysis of the spindle structure and chromosome distribution after IVM, a subgroup of oocytes from all groups were analyzed (n=20-25 oocytes per experimental group). After IVM culture, oocytes were fixed for 30 min at 37°C in a microtubule stabilizing buffer (MTSB-XF). A triple-labeling protocol was then used for the detection of microtubules, microfilaments and chromatin by fluorescence microscopy, as described previously (Messinger and Albertini, 1991; Costa-Borges et al., 2020). Briefly, fixed oocytes were first incubated with a mouse monoclonal against alpha-tubulin (LifeTechnologies-Invitrogen, 13-800), followed by an incubation with chicken anti-mouse IgG conjugated to Alexa Fluor 488 (Invitrogen, A21200) and Phalloidin staining for F-Actin conjugated with Alexa Fluor 594 (Invitrogen, A12381). Finally, oocytes were washed in PBS blocking solution, incubated in Hoechst 33258 (Millipore Sigma), and put on a mounting solution droplet on a glass slide. Stained oocytes were examined using an epifluorescence microscope (BX43F, Olympus) and digital images were acquired. Meiotic spindles with a bipolar barrel shape and chromosomes aligned in the metaphase II plate were considered as morphologically normal. Meiotic spindles with an abnormal (elongated bipolar or multipolar) shape and dispersed chromosomes were considered abnormal (Messinger and Albertini, 1991).

### ICSI and embryo culture

Mature M2 oocytes, in which the first polar body had completed its extrusion, were selected to be inseminated by ICSI. ICSI was performed with fresh mouse spermatozoa collected from the cauda epididymis of adult male mice (n=16), using a piezo drill-based protocol optimized for the mouse species (Costa-Borges et al., 2017). Briefly, fresh mouse sperm was collected and placed in a microdroplet of culture medium for 15 min at 37°C, 6% COD, and 7% OD concentration. Following incubation, 3 μl of the concentrated sperm solution was further diluted into a 150 μl droplet of culture medium, and ICSI was performed. Once injected, oocytes were washed thoroughly and cultured in groups of 5-8 embryos per drop in KSOM medium under mineral oil (Hypure oil Heavy, Kitazato) at 37°C, 6% COD, and 7% OD. The embryo culture was evaluated daily until Day 5 (96 hours) to assess fertilization, cleavage, and blastocyst rates.

### Embryo assessment through cell counts and immunofluorescence

On Day 5 of embryo culture, embryos that had progressed to the blastocyst stage were vitrified using Kitazato’s protocol and subsequently stored in liquid nitrogen, ready for embryo transfer on the next phase of the study (*in vivo* study, Figure 1).

A subset of embryos from each experimental group was cultured until Day 6 (120 hours) of development to conduct an additional validation aimed at assessing embryo quality (n=21-29 embryos per group). Differential cell counts were performed in the expanded blastocysts from each group to allow for a more objective evaluation of the trophectoderm (TE) and inner cell mass (ICM) cells. Briefly, blastocysts were fixed with 2% paraformaldehyde and subsequently permeabilized with 2.5% Triton X-100 overnight. On the next day, embryos were incubated with a mouse monoclonal antibody against Oct4 stem cell marker (sc-5279, Santa Cruz Biotechnology), followed by an incubation with a Alexa Fluor™ 488 donkey anti-mouse IgG secondary antibody (A-21202, Thermo Fisher Scientific). After, embryos were incubated in a 10 μg/ml bisbenzimide solution (Hoechst 33342, Millipore Sigma) for DNA staining. The fixed and stained blastocysts were then mounted on a microscope slide in a 1:1 PBS-glycerol droplet and flattened with a cover slip. Images were taken under epifluorescence microscopy (BX43F, Olympus). The total and ICM cell number were assessed manually with the assistance of an algorithm created by artificial intelligence (Costa-Borges et al., 2020).

### Embryo transfer and live birth analysis

On the day of the transfer, blastocysts vitrified from each group were warmed using Kitazato’s warming protocol, washed thoroughly, and kept in culture for 2 to 4 hours before embryo transfer. Surgical transfer procedures were performed using CD1 pseudopregnant females (n=56 mice). For each transfer, 4-5 embryos from each experimental condition were transferred per uterine horn. A total of 20 embryo transfer procedures were performed from embryos derived from each of the 4 study groups (Traditional IVM with denuded oocytes, OSC-IVM with denuded oocytes, Traditional IVM with COCs, and OSC-IVM with COCs), resulting in a total of 80 embryo transfers.

Following the embryo transfer procedures, recipients were allowed to deliver the first generation of offspring (F1). The resultant pups were raised by the surrogates until weaning. Pup weight and length were measured at Day 10 and Day 21, and sex ratio was determined on Day 10. Mouse behavior and development were monitored weekly to determine any abnormalities.

At reproductive age, five males and five females from each group were randomly selected and mated with wild type B6CBAF1 mice to produce the second offspring generation (F2). As in the F1, sex ratio, animal behavior, periodic weight, and global health status were monitored in the F2 generation and compared across groups.

### Determination of hormonal levels on F1 mice

To perform a hormonal assessment on serum, blood samples were collected from euthanized mice. Levels of follicle stimulating hormone (FSH), luteinizing hormone (LH), and progesterone (P4) were determined by immunochemiluminescence utilizing the Atellica IM Analyzer (Siemens Healthineers AG). Hormonal analysis in female mice was performed during the estrous stage of the reproductive cycle. To determine the stage of the estrous cycle vaginal smears were taken daily at 8 AM. Briefly, a pipette tip was inserted into the opening of the vagina with 70 μl of a normal saline solution. Several vaginal washes were performed with the pipette and the resulting fluid was collected with the pipette and transferred onto slides to perform a fresh microscopic examination. The estrous cycle phase of each animal was determined based on the cell types present: proestrus (mainly epithelial cells in the smear), estrus (high presence of squamous cells over epithelial cells and absence of leukocytes), and diestrus (leukocytes in fair abundance).

### Histological examinations on F1 mice

For histological assessment, tissue samples were obtained from F1 mice at the age of 4-6 months old from all four conditions: Traditional-IVM and OSC-IVM co-culture with both denuded oocytes and COCs. Specifically, the organs of five males and five females were analyzed per experimental group. In order to prepare mouse organs for collection, the animals were perfused with PBS followed by a 5% formaldehyde solution for tissue fixation. Afterward, tissues were collected and fixed overnight at 4°C in 5% formaldehyde. After fixation, tissues were embedded in paraffin wax, sectioned into 4 μm slices, and subjected to hematoxylin and eosin (H&E) staining. The heart, lungs, liver, kidneys, male or female reproductive tracts, and urinary bladder were then examined. Histological examination was conducted blindly using mouse identification codes for group assignments that were unknown to the evaluator.

### Statistical analysis

To ensure adequate statistical power for group comparisons, we determined that a total of 200 M2 oocytes per experimental group should undergo ICSI. To achieve this sample size, fourteen experimental replicates were performed for each study group. Results obtained from the experimental replicates were analyzed together. Statistical analyses were conducted utilizing Prism 10.0 software (GraphPad). The values of quantitative variables were analyzed using one-way analysis of variance (ANOVA) with follow up multiple comparison testing using the Sidak method between all experimental groups. P-values<0.05 were determined statistically significant.

## Results

We have previously demonstrated that *in vitro* maturation (IVM) of human immature cumulus oocyte complexes (COCs) using the OSC-IVM co-culture system significantly enhances the oocyte maturation rate and the formation of euploid blastocysts compared to traditional IVM media (Piechota et al., 2023). In this study, we aimed to perform a reprotoxicology study throughout multigenerational mouse development to further evaluate the safety of using human OSCs in the OSC-IVM system. For that, we leverage immature mouse oocytes, either denuded or enclosed in cumulus cells to model different oocyte dispositions after retrieval, and co-cultured them in OSC-IVM or in Traditional-IVM (Figure 2) to asses oocyte maturation, embryo development, and multigenerational health of F1 and F2.

**Figure 2.**
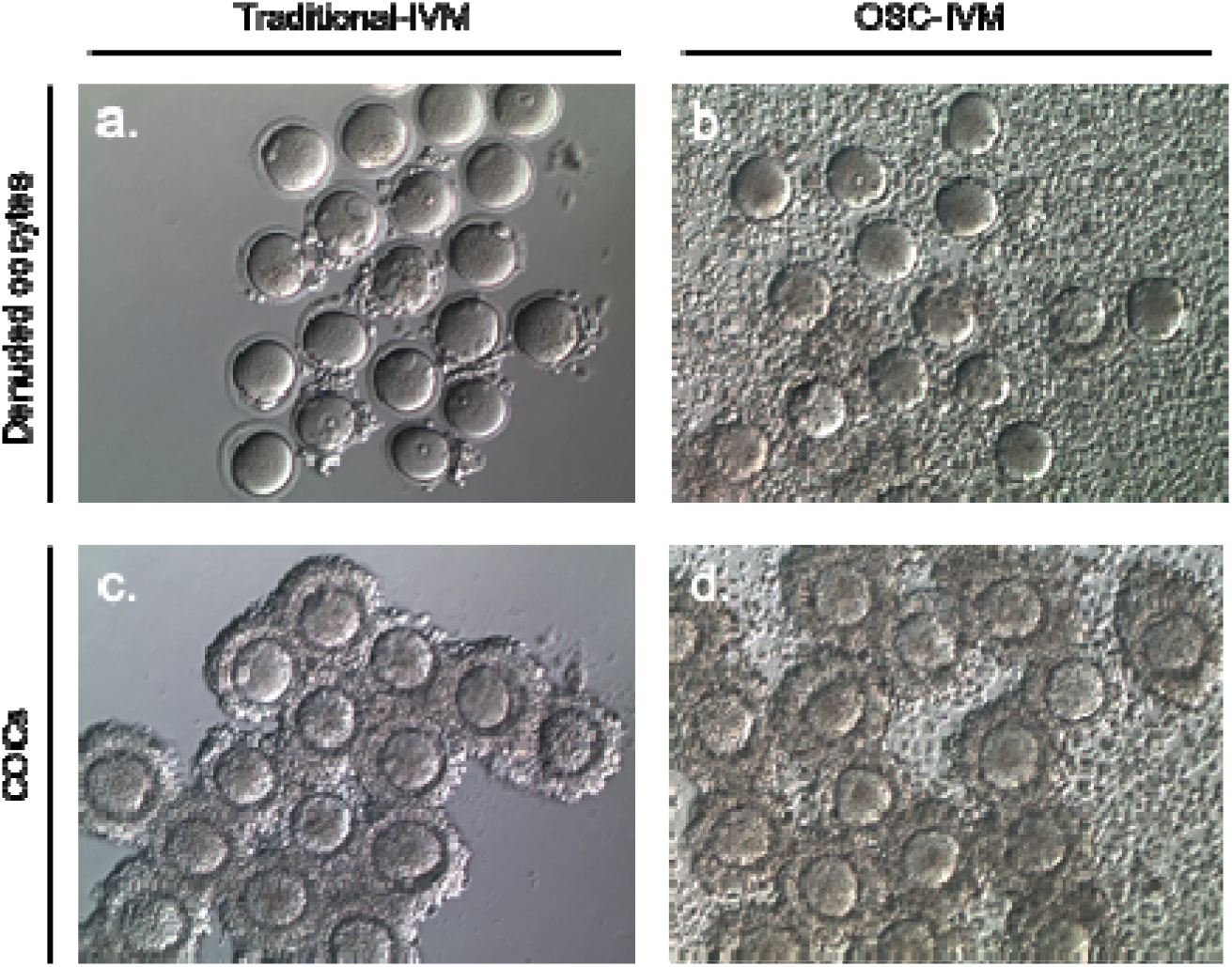
Representative images of murine oocyte *in vitro* maturation (IVM) using either OSC-IVM or Traditional-IVM culture systems. a. Denuded oocytes cultured in Traditional-IVM; b. Denuded oocytes cultured in OSC-IVM; c. Cumulus-oocyte complexes (COCs) cultured in Traditional-IVM; d. COC cultured in OSC-IVM.

### OSC-IVM improves blastocyst formation rate

The oocyte maturation rate of denuded oocytes and COCs cultured with the OSC-IVM system or Traditional-IVM was assessed after 18-20 hours of co-culture. A significant difference in oocyte maturation rate was found between the four groups (*p*=0.0099, ANOVA; Table 1). Unlike findings in humans, OSC-IVM generally resulted in a lower maturation rate of murine oocytes than Traditional-IVM, and the maturation rate of COCs was found to be lower than for denuded oocytes (Table 1). A significantly reduced maturation rate was observed in COCs cultured with OSC-IVM compared to Traditional-IVM with denuded oocytes (68.6% ± 14.1% versus 80.9% ± 5.9%, p=0.0101) (Table 1), while other groups did not differ significantly.

**Table 1.** Evaluation of *in vitro* oocyte maturation, fertilization and embryo development of OSC-IVM and Traditional IVM co-culture systems.

|  | Denuded oocytes |  | COCs |  | P value |
| --- | --- | --- | --- | --- | --- |
|  | Traditional-IVM<br>n= 466 | OSC-IVM<br>n= 464 | Traditional-IVM<br>n= 553 | OSC-IVM<br>n= 542 |  |
| Mature oocytes per total oocytes (%) mean ± SD | 80.9 ± 5.9 | 77.3 ± 9.4 | 78.5 ± 11.4 | 68.6 ± 14.1 | 0.0099<sup>a</sup> |
| Fertilized oocytes (%) per mature oocytes mean ± SD | 96.5 ± 4.8 | 95.0 ± 5.3 | 91.3 ± 5.2 | 89.5 ± 10.5 | 0.0055<sup>b</sup> |
| Cleavage embryos per fertilized oocytes (%) mean ± SD | 87.8 ± 9.8 | 89.1 ± 10.0 | 84.9 ± 12.5 | 88.3 ± 15.5 | 0.7547 |
| Blastocyst embryos per fertilized oocytes (%) mean ± SD | 38.0 ± 16.2 | 44.9 ± 18.5 | 41.5 ± 15.9 | 56.1 ± 19.2 | 0.0068<sup>c,d</sup> |

Abbreviations: COCs: Cumulus-Oocyte Complexes; N/A: Not applicable; SD: Standard deviation.
<sup>a</sup> Denuded oocytes Traditional-IVM vs. COCs OSC-IVM $p=0.0101$ , all others NS
<sup>b</sup> Denuded oocytes Traditional-IVM vs. COCs OSC-IVM $p=0.0098$ , All others NS
<sup>c</sup> Denuded oocytes Traditional-IVM vs. COCs OSC-IVM $p=0.0063$ , all other NS
<sup>d</sup> COCs Traditional-IVM vs. COCs OSC-IVM $p=0.0408$ , all others NS

Matured oocytes from both Traditional-IVM and OSC-IVM conditions were fertilized by ICSI, and we subsequently investigated the fertilization, cleavage, and blastocyst formation rates. A significant difference was found between fertilization rates of the four groups (p=0.0055, ANOVA; Table 1). Specifically, OSC-IVM with COCs significantly differed from Traditional-IVM with denuded oocytes (89.5 ± 10.5 versus 96.5 ± 4.8, p=0.0098), while there was no significant difference between other groups (Table 1). The assessment of cleavage-stage embryos during embryo culture revealed comparable cleavage rates between Traditional IVM and OSC-IVM treatments, with both groups yielding an 87.8-89.1% rate of embryo cleavage and no statistical differences among IVM conditions (p=0.7547, ANOVA; Table 1). Interestingly, a significant difference was found between the groups for blastocyst formation (p=0.0068, ANOVA; Table 1), with OSC-IVM demonstrating a higher formation rate than Traditional-IVM for both denuded oocytes and COCs. Specifically, blastocyst formation was statistically significant increased for the OSC-IVM treatment of COCs compared to Traditional IVM in COCs and denuded oocytes (56.1% ± 19.2% versus 41.5% ± 15.9% and 38.0% ± 16.2%; p=0.0408, and p=0.0063; Table 1), while no significant difference was observed between other groups.

### OSC-IVM leads to oocytes and embryos of similar morphological quality comparable to traditional *in vitro* matured oocytes

We then sought to compare the quality features of oocytes that underwent both IVM conditions, as well as the quality of the resulting blastocyst embryos after embryo culture. We firstly analyzed the status of the oocyte meiotic spindle by immunofluorescence, applying a triple staining protocol to study the chromosomal alignment and spindle assembly of M2 oocytes, which is a used methodology to assess oocyte quality (Coticchio et al., 2004). When we compared the spindle morphology among denuded oocytes and COCs cultured in Traditional-IVM or OSC-IVM conditions, we observed no differences between IVM groups (p=0.1136; Figure 3A-B). Spindle morphology analysis showed 100% normal spindle morphology on denuded oocytes cultured in both Traditional-IVM and OSC-IVM conditions, and 86-87% normal spindle morphology on COCs cultured in both IVM conditions (Figure 3A-B). Despite a limited number of oocytes used for this analysis, our findings suggest that OSC-IVM does not compromise oocyte quality.

**Figure 3.**
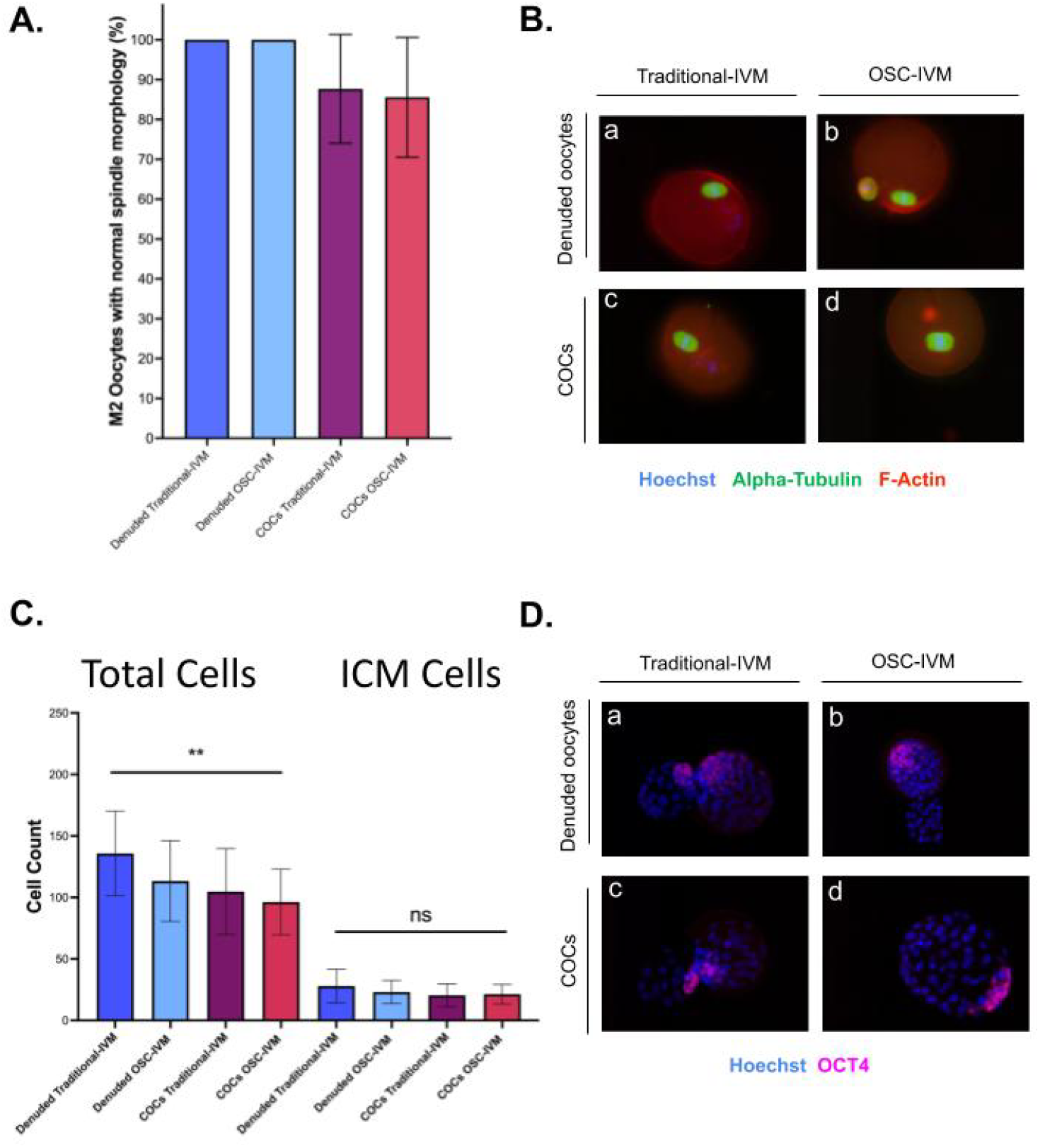
Assessment of oocyte and embryo quality by immunofluorescence. (A) Percentage of M2 oocytes with normal spindle morphology among denuded and cumulus-oocyte complexes (COCs) cultured in Traditional-IVM or OSC-IVM conditions (n=20-25 oocytes per group). (B) Representative immunofluorescence images of the meiotic spindles from each group used to analyze oocyte quality. Alpha-tubulin (microtubules) is stained in green, F-Actin (phalloidin) is stained in red, while chromosomes are stained with Hoechst in blue. (C) Total cells and inner cell mass (ICM) were assessed per embryo at D6 of embryo development (n=21-29 embryos per group). Error bars indicate mean ± SD. (D) Representative immunofluorescence images of blastocysts for each study group used to perform differential cell counts. The Oct4 germ cell marker (magenta) allows the identification of ICM, while the DNA marker Hoechst (blue) enables the identification of trophectoderm blastocyst cells. ^a^ Denuded oocytes Traditional-IVM vs. COCs OSC-IVM p=0.0030, all others NS.

To assess the potential impact of OSC-IVM on embryo development, we conducted an immunofluorescence analysis of embryos using the Oct4 germ cell marker, which allows for the identification of the ICM, along with the DNA marker Hoechst for trophectoderm blastocyst cells identification. As expected, no significant differences were observed in the inner cell count (p=0.1550, ANOVA) (Figure 3C-D). Interestingly, a significant difference in total cell count was observed between the groups (p=0.0047, ANOVA; Figure 3C-D) and the embryos from denuded oocytes had a generally higher number of total cells. The significant difference was found to be between OSC-IVM of COCs and Traditional-IVM of denuded oocytes, while no others significantly differed (p=0.0030, Figure 3C).

### Absence of reproductive toxicology in the F1 generation following OSC-IVM treatment

In order to evaluate the potential reproductive toxicological effects of OSC-IVM on live birth, blastocyst embryos generated using OSC-IVM and Traditional-IVM matured oocytes were vitrified and stored for subsequent embryo transfer into pseudopregnant mice. The blastocysts were warmed and prepared for embryo transfer in different replicates. No significant differences were observed in the morphology and quality of blastocysts derived from OSC-IVM and Traditional-IVM (Figure 4A). Importantly, all pregnant females delivered live pups, with no statistically significant differences observed between groups per birthing female, resulting in a mean of 28.2% to 34.3% of live births per transfer (p=0.6110, ANOVA; Table 2).

**Figure 4.**
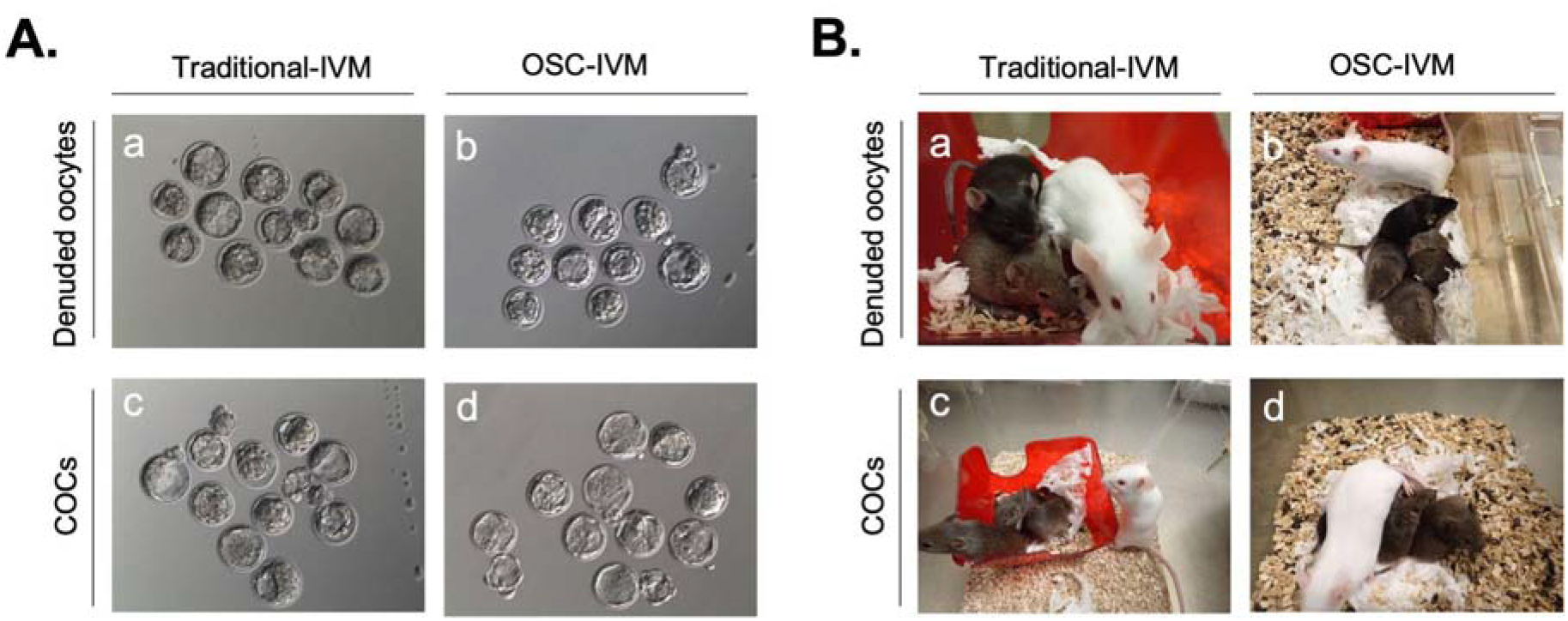
Representative images of murine blastocysts pre-embryo transfer (A) and subsequent live births (B) derived from oocytes matured *in vitro* using either OSC-IVM or Traditional-IVM co-culture systems. a. Denuded oocytes cultured in Traditional-IVM; b. Denuded oocytes cultured in OSC-IVM; c. Cumulus-oocyte complexes (COCs) cultured in Traditional-IVM; d. COCs cultured in OSC-IVM.

**Table 2.** Analysis of live births per birthing female.

|  | Denuded oocytes |  | COCs |  | P value |
| --- | --- | --- | --- | --- | --- |
|  | Traditional-IVM | OSC-IVM | Traditional-IVM | OSC-IVM |  |
| **F1 Live birth per transfer (%) mean ± SD** | 28.2 ± 19.4 | 29.0 ± 0.07 | 34.3 ± 17.3 | 25.3 ± 13.0 | 0.6110 |
| **F1 Day 10 Pup weight (grams) mean ± SD** | 8.8 ± 1.6 | 9.2 ± 1.2 | 9.4 ± 1.2 | 9.7 ± 2.2 | 0.2444 |
| **F1 Day 10 Pup length (cm) mean ± SD** | 5.5 ± 1.2 | 5.0 ± 0.3 | 5.1 ± 0.4 | 5.2 ± 0.7 | 0.1251 |
| **F1 Sex Ratio (% males) mean ± SD** | 45.8 ± 29.7 | 49.0 ± 32.7 | 46.9 ± 37.2 | 54.2 ± 40.0 | 0.9751 |
| **F1 Day 21 Pup weight (grams) mean ± SD** | 15.9 ± 1.2 | 16.6 ± 2.2 | 16.1 ± 1.9 | 16.1 ± 3.72 | 0.5976 |
| **F1 Day 21 Pup length (cm) mean ± SD** | 6.8 ± 0.3 | 7.0 ± 0.4 | 7.0 ± 0.4 | 7.0 ± 0.7 | 0.2135 |
| **F1 Developmental or Behavioral Abnormalities** | None noted | None noted | None noted | None Noted | N/A |
| **F1 Fecundity (F2 pups born per F1 outbreeding) (N) mean ± SD** | 8.2 ± 1.5 | 6.8 ± 1.1 | 7.6 ± 2.2 | 8.2 ± 1.6 | 0.2040 |
| **F2 Sex Ratio (% males) mean ± SD** | 53.9 ± 20.0 | 49.3 ± 10.0 | 52.2 ± 20.0 | 51.5 ± 20.0 | 0.9608 |
| **F2 Developmental or Behavioral Abnormalities** | None noted | None noted | None noted | None Noted | N/A |

Abbreviations: COCs: Cumulus-oocyte complexes; F1: first generation; F2: second generation; N/A: Not applicable; SD: Standard deviation.

When the F1 generation was analyzed, no significant differences were observed in the pup weight and length at either Day 10 and Day 21 of development between the different conditions (Table 2). Furthermore, there were no significant differences in the number of males/females born per group, and no developmental or behavioral abnormalities were reported on the F1 generation (p=0.9751, ANOVA; Figure 4B, Table 2). Additionally, mice were monitored weekly according to standard patterns of mice welfare until reaching adulthood, further ensuring no physiological or behavioral alterations. Additionally, no signs of sickness or disease were reported for any of the females (F0) who received an embryo transfer from embryos derived from either OSC-IVM or Traditional-IVM conditions.

Moreover, to further ensure the absence of long-term implications and transgenerational effects derived from OSC-IVM culture, we evaluated the fecundity of the F1 generation and the resulting health and development of the F2 generation. No significant differences were found in fecundity of F1 mice between groups, as all groups yielded a similar range of offspring, with litter sizes ranging from 6.8 to 8.2 pups (p=0.2040, ANOVA; Table 2). Furthermore, no differences were noted in the sex ratio of F2 generation, and no developmental or behavioral abnormalities were reported (Table 2). Overall, the result from this reprotoxicology study suggests that OSC-IVM does not appear to have any significant impact on reproduction, progeny health, or progeny fecundity.

### Assessment of F1 reproductive function through hormonal evaluation

To further confirm the absence of reproductive alterations in the progeny, the serum values of reproductive-related hormones were also analyzed on the F1 generation (Table 3). Mice from the different experimental groups showed regular serum values of FSH, LH, and P4, with no statistically significant differences between conditions for males and females, confirming the maintenance of a normal reproductive function across all experimental conditions.

**Table 3.** Serum values of reproductive-related hormones on F1 generation.

|  | Denuded oocytes |  | COCs |  | P value |
| --- | --- | --- | --- | --- | --- |
|  | Traditional-IVM | OSC-IVM | Traditional-IVM | OSC-IVM |  |
| Males |  |  |  |  |  |
| FSH (IU/L) | 2.0 ± 0.5 | 1.9 ± 0.5 | 1.6 ± 0.8 | 1.4 ± 0.4 | 0.4188 |
| LH (IU/L) | 1.2 ±1.2 | 0.6 ± 0.1 | 0.6 ± 0.2 | 0.6 ± 0.3 | 0.3340 |
| P4 (ng/mL) | 7.9 ± 1.8 | 4.3 ± 1.2 | 8.7 ± 5.9 | 10.8 ± 6.9 | 0.2173 |
| Females |  |  |  |  |  |
| FSH (IU/L) | 1.5 ± 0.3 | 1.8 ± 0.4 | 1.8 ± 0.6 | 1.9 ± 0.3 | 0.4393 |
| LH (IU/L) | 0.7 ± 0.2 | 0.6 ± 0.2 | 0.8 ± 0.4 | 0.9 ± 0.2 | 0.4050 |
| P4 (ng/mL) | 14.0 ± 5.3 | 13.0 ± 7.3 | 22.2 ± 20.4 | 6.9 ± 4.6 | 0.2427 |

Abbreviations: COCs: Cumulus-oocyte complexes; FSH: Follicle-stimulating hormone; LH: Luteinizing hormone; P4: Progesterone. Data are presented as mean ± standard deviation.

### OSC-IVM does not lead to any tissue pathological anomalies

To further explore overall health and well-being of the progeny, we then performed a comprehensive histopathological evaluation on F1 generation mice for all experimental conditions. Morphological analyses on H&E stained tissue were performed on the main organs, including the brain, heart, lungs, liver, kidney, urinary bladder, as well as the male and female reproductive organs. Despite a limited number of animals being assessed per group (n=4-5 per condition), our findings revealed no specific pathological abnormalities associated with any particular experimental group (Table 4, Suppl. Fig. 1), providing additional evidence demonstrating the absence of adverse effects attributable to OSC-IVM treatment.

**Table 4.** Histopathology assessment of F1 generation mice. See representative H&E images of normal and altered histology for each organ in Supplementary Figure 1.

|  | Males |  |  |  | Females |  |  |  |
| --- | --- | --- | --- | --- | --- | --- | --- | --- |
|  | Denuded oocytes |  | COCs |  | Denuded oocytes |  | COCs |  |
|  | Traditional IVM | OSC-IVM | Traditional IVM | OSC-IVM | Traditional IVM | OSC-IVM | Traditional IVM | OSC-IVM |
| **Brain** |  |  |  |  |  |  |  |  |
| Normal histology | 4/5 | 5/5 | 5/5 | 4/4 | 5/5 | 5/5 | 5/5 | 5/5 |
| Vacuoles in hippocampal neurons | 1/5 | 0/5 | 0/5 | 0/5 | 0/5 | 0/5 | 0/5 | 0/5 |
| **Heart** |  |  |  |  |  |  |  |  |
| Normal histology | 5/5 | 4/5 | 5/5 | 4/4 | 5/5 | 5/5 | 5/5 | 5/5 |
| inflammatory infiltration (1) | 0/5 | 1/5 | 0/5 | 0/4 | 0/5 | 0/5 | 0/5 | 0/5 |
| **Lungs** |  |  |  |  |  |  |  |  |
| Normal histology | 5/5 | 5/5 | 5/5 | 4/4 | 4/5 | 5/5 | 4/5 | 4/5 |
| inflammatory infiltration (2) | 0/5 | 0/5 | 0/5 | 0/4 | 1/5 | 0/5 | 1/5 | 1/5 |
| **Liver** |  |  |  |  |  |  |  |  |
| Normal histology | 4/5 | 3/5 | 1/5 | 0/4 | 4/5 | 2/5 | 2/5 | 3/5 |
| Glycogen deposition | 1/5 | 2/5 | 4/5 | 4/4 | 0/5 | 0/5 | 2/5 | 2/5 |
| inflammatory infiltration (3) | 0/5 | 0/5 | 0/5 | 0/5 | 0/5 | 3/5 | 1/5 | 0/5 |
| Areas of hepatocyte necrosis | 0/5 | 0/5 | 0/5 | 0/5 | 1/5 | 0/5 | 0/5 | 0/5 |
| **Kidney** |  |  |  |  |  |  |  |  |
| Normal histology | 2/5 | 2/5 | 2/5 | 1/4 | 4/5 | 4/5 | 4/5 | 3/5 |
| Cytoplasmic vacuolation | 2/5 | 1/5 | 2/5 | 1/4 | 0/5 | 0/5 | 0/5 | 0/5 |
| inflammatory infiltration (4) | 1/5 | 2/5 | 1/5 | 2/4 | 1/5 | 1/5 | 1/5 | 2/5 |
| **Urinary bladder** |  |  |  |  |  |  |  |  |
| Normal histology | 5/5 | 4/5 | 5/5 | 3/4 | 4/5 | 2/5 | 2/5 | 3/3 |
| inflammatory infiltration (2) | 0/5 | 1/5 | 0/5 | 1/4 | 1/5 | 3/5 | 3/5 | 0/3 |
| **Reproductive organs** |  |  |  |  |  |  |  |  |
| **Testis** |  |  |  |  |  |  |  |  |
| Normal histology | 1/5 | 1/5 | 1/5 | 0/4 | - | - | - | - |
| Initial testicular degradation | 2/5 | 4/5 | 2/5 | 3/4 | - | - | - | - |
| Severe testicular degradation (5) | 2/5 | 0/5 | 2/5 | 1/4 | - | - | - | - |
| **Epididymis** |  |  |  |  |  |  |  |  |
| **Normal histology** | 5/5 | 5/5 | 4/5 | 4/4 | - | - | - | - |
| **Obstruction of the epididymis** | 0/5 | 0/5 | 1/5 | 0/4 | - | - | - | - |
| **Ovary** |  |  |  |  |  |  |  |  |
| **Normal histology** | - | - | - | - | 4/4 | 5/5 | 4/5 | 4/5 |
| **Follicular cyst** | - | - | - | - | 0/4 | 0/5 | 1/5 | 1/5 |
| **Uterus** |  |  |  |  |  |  |  |  |
| **Normal histology** | - | - | - | - | 3/5 | 4/5 | 4/5 | 2/5 |
| **Stromal oedema** | - | - | - | - | 2/5 | 1/5 | 1/5 | 1/5 |
| **Cystic endometrial hyperplasia** | - | - | - | - | 0/5 | 0/5 | 0/5 | 1/5 |
| **inflammatory infiltration (6)** |  |  |  |  | 0/5 | 0/5 | 0/5 | 1/5 |
| **Haemorrhagic area** | - | - | - | - | 0/5 | 0/5 | 0/5 | 0/5 |

(1) Inflammatory infiltration predominantly consisting of polymorphonuclear neutrophils in the aortic tunica media and adventitia.
(2) Inflammatory infiltration as indicated by the presence of a lymphoplasmacytic infiltrate.
(3) Focus of inflammation composed of lymphocytes and polymorphonuclear neutrophils.
(4) Focus of lymphoplasmacytic infiltrate observed around arcuate vessels.
(5) Includes cases of reduced number or depletion of germ cells, along with degeneration of seminiferous tubules.
(6) Focus of inflammation composed of macrophages and polymorphonuclear neutrophils.

## Discussion

*In vitro* maturation (IVM) is an emerging technique that involves the retrieval of immature oocytes from developing antral follicles from minimally stimulated or unstimulated ovaries, followed by their maturation in a laboratory setting prior to their utilization for IVF purposes or egg freezing procedures. While current IVM methodologies have demonstrated success in yielding mature oocytes and facilitating embryo formation, pregnancies and healthy live births (Akin et al., 2021; Braam et al., 2019; De Vos et al., 2011; Fadini et al., 2009; Guzman et al., 2012; Ma et al., 2020; Mochini et al., 2011; Mohsenzadeh et al., 2022; Piechota et al., 2023; Pongsuthirak et al., 2015; Sanchez et al., 2019; Sanchez et al., 2017; Shu-Chi et al., 2006; Vuong et al., 2020a, 2020b), their rates are lower compared to conventional IVF, highlighting the need for more research aimed at developing innovative approaches for improving IVM outcomes by optimizing oocyte maturation conditions and enhancing oocyte quality.

Our group has recently developed a pioneering IVM system based on the use of hiPSC-derived OSC to recapitulate the dynamic ovarian function *in vitro*, known as OSC-IVM (Piechota et al., 2023; Pierson Smela et al., 2023). In our previous studies, we demonstrated that co-culture of immature human COCs from minimal stimulation cycles in OSC-IVM system significantly enhanced oocyte maturation rate and improved euploid day 5 or 6 blastocyst formation rates, compared to a commercially available Traditional IVM system (Piechota et al., 2023). By enhancing the maturation of oocytes *in vitro*, OSC-IVM has emerged as a promising tool for assisted reproduction, highlighting the potential of cell engineering approaches for improving ART treatments and patient outcomes.

Notably, recent findings demonstrate that OSC-IVM can also be applied to improve the maturation rate of human denuded oocytes from conventional stimulation cycles, a process known as rescue IVM (Giovannini et al., 2023). The results of the current study show that OSC-IVM improves blastocyst formation compared to Traditional-IVM, and further that cumulus enclosure prior to IVM improves blastocyst formation beyond IVM in denuded oocytes, in line with previous findings. These results suggest that OSC-IVM likely improves oocyte cytoplasmic maturation via paracrine factor exchange, primarily via interaction with cumulus cells and to a lesser extent oocytes themselves. Therefore, these findings support that the most efficacious performance of OSC-IVM is with COCs, but that OSC-IVM can still improve developmental outcomes even in denuded oocytes. A potential model for this finding is that OSC-IVM recapitulates a follicle-like environment by establishment and maintenance of the hormonal, growth factor, and steroid environment of ovarian follicles *in vitro* and dynamic communication between oocytes, cumulus cells and OSCs (Supplementary Figure 2).

The present study provides comprehensive examination of the safety of OSC-IVM, employing a multifaceted approach including *in vitro* and *in vivo* reproductive assessments, behavioral evaluations and analyses of hormonal and histological parameters. Overall, our study included more than 2000 immature denuded oocytes and cumulus-enclosed oocytes, allowing for a robust comparison of the safety profile of OSC-IVM compared to a commercially available option for traditional IVM. During the initial *in vitro* phase, we demonstrated an improvement on the embryo blastocyst rates of murine COCs and denuded oocytes cultured in OSC-IVM system, compared to a traditional IVM media, aligning with our previous observations in humans (Piechota et al., 2023; Giovannini et al., 2023).

However, unlike in humans, OSC-IVM does not improve oocyte maturation in mice. This disparity may originate from interspecies variability, particularly in the utilization of human OSCs to stimulate a response in mouse oocytes. In fact, prior research has highlighted distinct molecular mechanisms underlying oocyte maturation between human and murine species (Zhao et al., 2020). Despite the primary focus of this study being to assess potential adverse effects of OSC-IVM on embryo development and subsequent offspring, our findings are intriguing, as they suggest that OSC-IVM may actually enhance oocyte cytoplasmic maturation despite interspecies differences. This improvement could potentially explain the observed increase in the blastocyst formation rate.

Nevertheless, the analysis of the spindle morphology, known to be a marker of oocyte quality (Coticchio et al., 2004), revealed no differences in the spindle structure of M2 oocytes between OSC-IVM and Traditional-IVM. Additionally, these maturated oocytes yielded high-quality embryos with no differences in the total embryo cell counts, ICM cells counts or *in vivo* developmental rates between groups. Therefore, these findings suggest that OSC-IVM maintains oocyte quality and does not compromise early embryo development or live birth rates, indicating its suitability for ART procedures.

Furthermore, this multigenerational follow-up of progeny birth, health, development, and fecundity demonstrates that compared to Traditional-IVM, OSC-IVM does not display any evidence of reproductive toxicity. Both the progeny derived from OSC-IVM-treated oocytes, with and without cumulus cell enclosure, and the recipient females who received embryos from OSC-IVM-treated oocytes displayed no significant adverse health side effects or developmental abnormalities attributed to the treatment. Particularly given that OSC-IVM leverages allogeneic hiPSC-derived somatic cell co-culture with gametes, this study importantly demonstrates that no detectable or lasting effects of OSC-IVM treatment were observed or any evidence of residual cell activity or presence beyond the oocyte maturation stage. These findings corroborate our previous preclinical study showing high-quality oocyte and embryo development in humans arising from OSC-IVM treatment (Piechota et al. 2023) as well as similar studies in the field that have leveraged primary granulosa cell co-cultures to improve IVM outcomes. Overall, this study suggests that OSC-IVM is a generally effective IVM approach that does not display apparent reproductive toxicity, which is an important finding to support its future clinical investigation.

### Limitations and Reasons for Caution

Although a high number of oocytes were analyzed in this study, enough to achieve statistical results in the variables studied, the number of oocytes and embryos available were lower for some of the techniques performed, particularly the oocyte spindle analysis and embryo cell count analysis. A higher number of oocytes and embryos could be helpful to perform the fluorescence analysis, confirming the results achieved.

This study was conducted in compliance with European Medicines Agency (EMA) ICH E6 (R2) Good clinical practice scientific guidelines to demonstrate the OSC-IVM safety. Nevertheless, human clinical studies evaluating live births are necessary to ensure the safety of the stem cell-derived OSCs platform for IVM. In general, the inherent species specific differences between the murine oocytes and human OSCs tested here represents a limit of common reproductive toxicology assessment approaches and necessitates further studies in humans.

## Acknowledgements

This work was carried out with the support of Embryotools S.L, Barcelona, Spain. We thank the dedicated support and work of the Embryotools team for coordinating and managing this collaborative study. We additionally thank the Animal facility of the Science Park of Barcelona for their contribution to performing animal procedures.

## Authors’ roles

C.C.K. and N.C-B. designed, supervised and coordinated the study; E.M., M.A., N.C-B., performed the research study, F.B., K.S.P., S.P., B.P., A.D.N., produced and qualified OSC batches, M.M., F.B., C.C.K., performed data interpretation and statistical analysis, M.M., K.S.P., A.B.F., coordinated study logistics, M.M., F.B., C.C.K. wrote the manuscript with significant input from all authors.

## Data availability

All data needed to evaluate the conclusions in the paper are present in the paper and supplementary tables and figures. Anonymized raw data for all findings in the paper will be provided upon request.

## Supplementary Figures

**Supplementary Figure 1.**
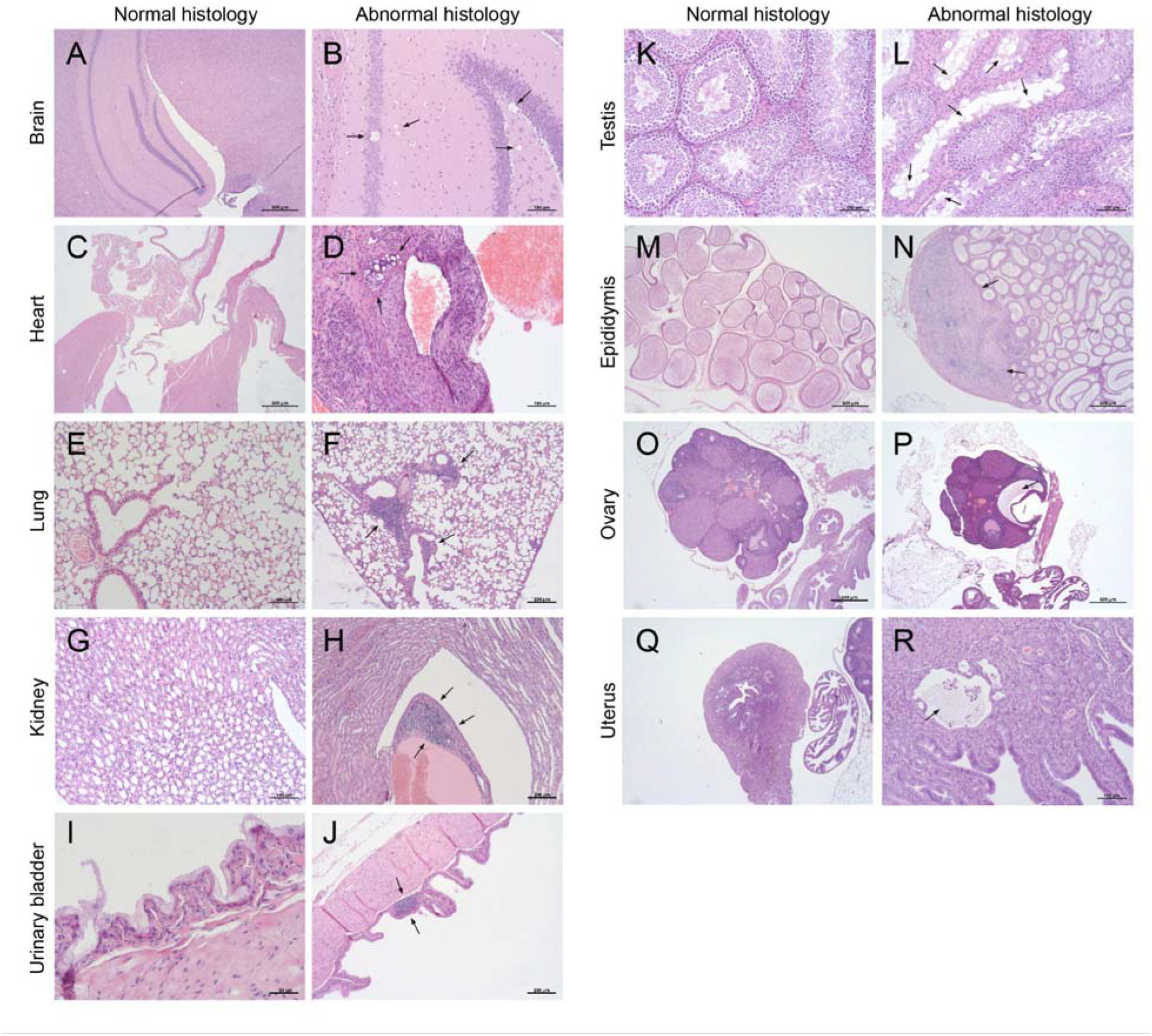
Representative images of the histological evaluation of organs in F1 generation mice. H&E staining shows representative images of tissues with normal histology (left panels) compared to tissues with altered histology (right panels) for each of the organs evaluated. A) Normal brain hippocampus, B) Brain hippocampus with presence of some vacuoles in hippocampal neurons (arrows), C) Normal heart displaying aorta (A), left atrium (LA), aortic valve (AV), mitral valve (MV), right ventricle (RV) and left ventricle (LV), D) Presence of cartilaginous tissue in the base of the heart (arrows) accompanied by inflammatory infiltration in the aortic wall, E) Normal lung, F) Lung with presence of a focal peribronchiolar lymphoplasmacytic infiltration (arrows), G) Normal kidney medulla, H) Kidney with lymphoplasmacytic infiltrate around renal arcuate vessels (arrows), I) Normal urinary bladder, J) Urinary bladder with presence of lymphoplasmacytic infiltrate in lamina propria (arrows), K) Normal testicle showing presence of complete spermatogenesis within the seminiferous tubules, L) Testicle exhibiting testicular degeneration, characterized by scattered seminiferous tubules containing vacuolated spermatogenic cells and a reduced number of male germ cells (arrows), M) Normal epididymis, N) Epididymis exhibiting aspermia secondary to tubule obstruction (arrows) and macrophage infiltration, O) Ovary with primary, secondary, tertiary follicles and corpus luteum, P) Ovary with presence of follicular cyst (arrow), Q) Normal uterus, and R) Uterus with a dilated gland containing macrophages, polymorphonuclear neutrophils, and cellular debris (arrow). The scale bars indicate the magnitude for each individual image.

**Supplementary Figure 2:**
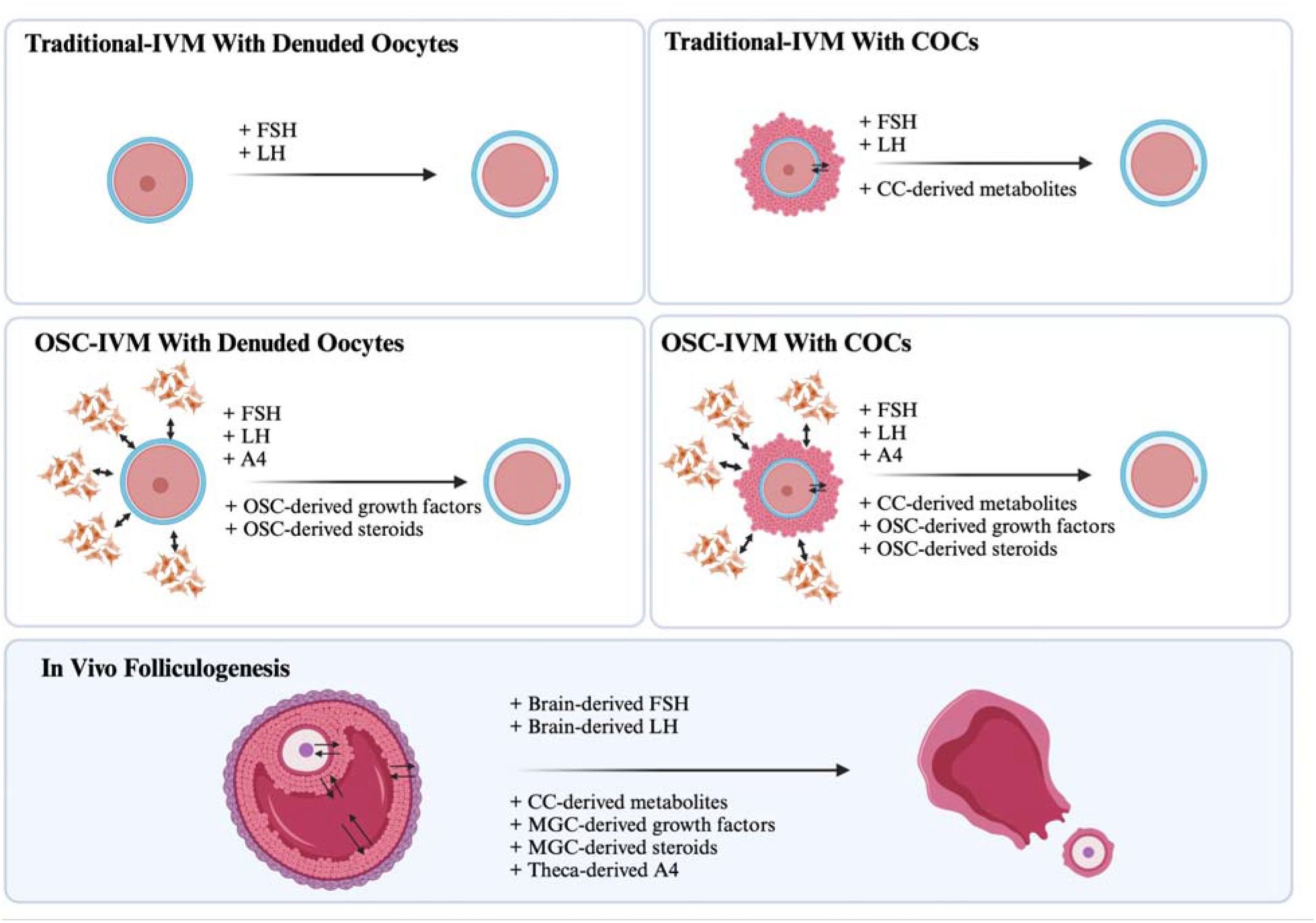
Model of mechanism of action of OSC-IVM compared to Traditional-IVM and *in vivo* maturation. Abbreviations: follicle stimulating hormone, FSH; luteinizing hormone, LH; cumulus cell, CC; ovarian support cells, OSC; androstenedione, A4; mural granulosa cells, MGC; cumulus-oocyte complexes, COCs.

